# CD44 inhibits α-SMA gene expression via a novel G-actin/MRTF mediated pathway that requires TGFβR/p38MAPK activity in murine skin fibroblasts

**DOI:** 10.1101/676635

**Authors:** Yan Wang, Judith A. Mack, Edward V. Maytin

## Abstract

Well-regulated differentiation of fibroblasts into myofibroblasts (MF) is critical for skin wound healing. Neoexpression of α-smooth muscle actin (α-SMA), an established marker for MF differentiation, is driven by TGFβ receptor (TGFβR)-mediated signaling. Hyaluronan (HA) and its receptor CD44 may also participate in this process. To further understand this process, primary mouse skin fibroblasts were isolated and treated *in vitro* with recombinant TGF-β1 (rTGF-β1) to induce α-SMA expression. CD44 expression was also increased. Paradoxically, CD44 knockdown by RNA interference (RNAi) led to increased α-SMA expression and α-SMA-containing stress fibers. Removal of extracellular HA or inhibition of HA synthesis had no effect on α-SMA levels, suggesting a dispensable role for HA. Exploration of mechanisms linking CD44 knockdown to α-SMA induction, using RNAi and chemical inhibitors, revealed a requirement for non-canonical TGFβR signaling through p38MAPK. Decreased monomeric G-actin but increased filamentous F-actin following CD44 RNAi suggested a possible role for Myocardin-related Transcription Factor (MRTF), a known regulator of α-SMA transcription and itself being regulated by G-actin binding. CD44 RNAi promoted nuclear accumulation of MRTF and the binding to its transcriptional cofactor, SRF. MRTF knockdown abrogated the increased α-SMA expression caused by CD44 RNAi, suggesting that MRTF is required for CD44-mediated regulation of α-SMA. Finally, chemical inhibition of p38MAPK reversed nuclear MRTF accumulation after rTGF-β1 addition or CD44 RNAi, revealing a central requirement for p38MAPK in both cases. We concluded that CD44 regulates α-SMA gene expression through cooperation between two intersecting signaling pathways, one mediated by G-actin/MRTF and the other via TGFβR/p38MAPK.

## INTRODUCTION

Fibrosis is a common process that involves differentiation of fibroblasts into myofibroblasts (MF), which are cells that that mediate cellular contraction and extracellular matrix production. Fibrosis can be a normal and necessary process during tissue repair (e.g., scar formation after cutaneous wounding) (1,2). However, dysregulated fibrosis gives rise to many pathological conditions including keloids(3), scleroderma (4), hepatic cirrhosis(5), and idiopathic pulmonary fibrosis(6). Nascent and cumulative expression of α-smooth muscle actin (α-SMA) is one of the most well-established hallmarks for MF differentiation(7). Once generated, α-SMA can incorporate into actin stress fibers and produce contractile forces which play a critical role in wound closure and scar formation during wound healing(8). Multiple transcriptional regulatory proteins have been identified that respond to biomechanical and metabolic signals generated after skin injury; together with specific binding partners, these proteins bind to the α-SMA promoter and regulate α-SMA gene expression (9). One of these is serum response factor (SRF), a mammalian transcription factor that binds to a consensus sequence CArG box (CC(A/T)6GG). This serum response element (SRE) is found in smooth muscle cell-specific contractile genes(9). Myocardin-related transcription factor A (MRTF-A), a co-activator of SRF, is normally sequestered in the cytoplasm as a result of its binding to monomeric G-actin (globular actin). In response to growth factor stimulation, or after activation of Rho/ROCK signaling and subsequent polymerization of G-actin to F-actin (filamentous actin), the amount of G-actin is decreased and MRTF is freed from sequestration, allowing it to translocate into the nucleus and bind to SRF, hence triggering SRE-dependent α-SMA gene expression(10,11).

TGF-β and its downstream effectors constitute one of the most potent regulatory cascades for α-SMA gene expression and MF differentiation. Among the three TGFβ ligands (TGFβ1, 2, 3), TGFβ1 is the predominant isoform and also the most studied in the context of MF differentiation (12,13). Binding of TGF-β1 to TGFβ Receptor II (TβRII) facilitates the formation of a heterotetrameric complex between the TβRII homodimer and the TGFβ Receptor I (TβRI) homodimer. This results in the phosphorylation of the GS domain of TβRI and the activation of TβRI receptor kinase activity, leading to recruitment and phosphorylation of receptor-regulated Smads (R-Smads: Smad2 and Smad3). Phosphorylated Smad2 or Smad3 rapidly dissociate from TβRI and form a complex with Smad4, which translocates to the nucleus and activates α-SMA gene transcription(14). Although the Smad-dependent signaling pathway is widely accepted as the canonical pathway induced by TGF-β1, accumulating evidence has shown that TGF-β1 can also activate a number of other Smad-independent signaling pathways that involve mitogen-activated protein kinases (MAPKs) such as ERK1/2, JNK, p38MAPK, PI3K/AKT, and RhoA (15). Many of these molecules are also involved in α-SMA gene regulation in a tissue and cell contextdependent manner (15). In particular, studies from different groups have shown a role for p38MAPK signaling during fibrotic processes within tissues and organs as diverse as the kidney (16), eye (17) and cardiac muscle (18).

CD44 is a type I transmembrane proteoglycan with multiple isoforms that arise from alternative splicing of ten variable exons; the most widely expressed isoform is the standard form (CD44s)(19). CD44 not only mediates cell-cell and cell-matrix interactions, but also binds growth factors and interacts with a number of growth factor receptors within the cell membrane, thereby exerting a complex role in the regulation of cell growth, survival and differentiation(19-21). CD44 is also the principal receptor for hyaluronan (HA), a major constituent of the extracellular matrix and a mediator of some aspects of CD44 signaling (22). HA is a negatively charged, unbranched, non-sulfated glycosaminoglycan consisting of alternating disaccharide units of D-glucuronic acid and N-acetyl-glucosamine. HA is synthesized by HA-synthase enzymes (Has1, Has2, Has3) and degraded by hyaluronidases (Hyal1 and Hyal2)(23). Importantly, HA has been demonstrated to be involved in pro-fibrotic signaling and myofibroblast differentiation in human lung fibroblasts (24,25) and in the differentiation of epithelial cells and tissues as well (26,27).

When considering CD44 and HA and their role in regulating TGF-β-mediated profibrotic signaling and MF differentiation, many issues remain unresolved since different groups have generated contradictory results. In some experimental systems, both HA and CD44 appear to be supportive and stimulatory for TGFβ-driven MF differentiation. For example, Simpson et al showed that in human dermal fibroblasts, CD44 gene silencing inhibited TGF-β-induced α-SMA expression. They further demonstrated that TGF-β1-mediated MF differentiation required an interaction between EGFR, CD44, and HA in the plasma membrane which could be disrupted by treatment with hyaluronidase or 4-MU (28,29). As another example of a stimulatory role of HA and CD44 upon fibrosis and α-SMA expression, Li et al demonstrated that targeted overexpression of HAS2 in myofibroblasts generated an aggressive profibrotic phenotype in a bleomycin-induced model of lung fibrosis, that could be inhibited by CD44 depletion, or by blockade with a CD44 blocking antibody (30). On the other hand, evidence from other groups showed that CD44 can have an inhibitory effect on TGF-β receptor signaling and fibrosis in some tissues. Ansorge et al found that in a tendon-healing model, loss of CD44 (CD44 knockout mice) led to significant increases in the gene expression of TGF-β1, TGF-β3, and COL3A1, as well as in parameters indicative of good tissue repair such as material strength and tissue organization (31). Porsh et al (32) showed that CD44 interacts physically with TGFβR (and also platelet derived growth factor receptor, PDGFRβ) in the plasma membrane of primary human dermal fibroblasts. Depletion of CD44 promoted TGFβ receptor signaling activity, implying that CD44 is a negative regulator of TGFβ signaling in these fibroblasts. In a third example, Velasco et al (33) showed that knockout of CD44 stimulated a robust activation of TGFβ receptor II and collagen type III expression and improved dermal regeneration and wound closure in Adamts5-/- mice. These contrasting examples suggest that the mechanistic relationship between CD44 and TGFβ-mediated regulation of fibrotic responses remains unclear.

In this study, we sought to obtain a better understanding of how CD44 regulates TGF-β signaling and expression of α-SMA in primary murine dermal fibroblasts. We studied the impact of inhibiting CD44 gene expression by RNA interference and observed an upregulation of α-SMA. This was accompanied by changes in the cytoskeleton including an increase in F-actin polymerization, in parallel with an observed increase in nuclear accumulation of MRTF. Interestingly, CD44-mediated effects appeared to be entirely independent of HA, and although TGF-β signaling was also involved, p38MAPK rather than Smad2 turned out to be the relevant downstream effector.

## RESULTS

### CD44s is the predominant CD44 isoform in primary murine dermal fibroblasts and is inhibitory to α-SMA gene expression

The various isoforms of CD44, including the standard form (CD44s) and other variants, are expressed in a tissue and cell context-dependent manner (19). We examined the expression of CD44 in murine dermal fibroblasts, and in primary murine keratinocytes and epithelial cancer cells for comparison, on western blots using a polyclonal pan-CD44 antibody. As shown in Figure 1A, *left*, CD44s was the only detectable form of CD44 in fibroblasts, as confirmed by a specific knockdown experiment in which the other bands are not responsive to CD44 RNAi and therefore are nonspecific (Fig.1A, *right*), and in contrast to primary keratinocytes and MDA-MB-468 breast cancer cells that express CD44E (34) and CD44v6 (35), respectively. In primary murine fibroblasts, CD44 was shown to be upregulated at both the protein and mRNA level in response to treatment with rTGF-β1 (Figure 1B). Because CD44 has been shown to facilitate TGF-β-driven myofibroblast differentiation in human lung fibroblasts (28,36), we anticipated that increased CD44 gene expression after TGFβ treatment might contribute to the observed up-regulation of α-SMA. We tested this hypothesis by knocking down CD44 via RNA interference (RNAi). To our surprise, the expression of α-SMA was further increased in response to CD44 RNAi, at both the mRNA (Figure 1C) and protein levels (Figure 1D). In addition, the formation of α-SMA-positive stress fibers was significantly enhanced in fibroblasts treated with CD44 RNAi, as analyzed by immunofluorescent staining (Figure 1E).

**Figure 1.**
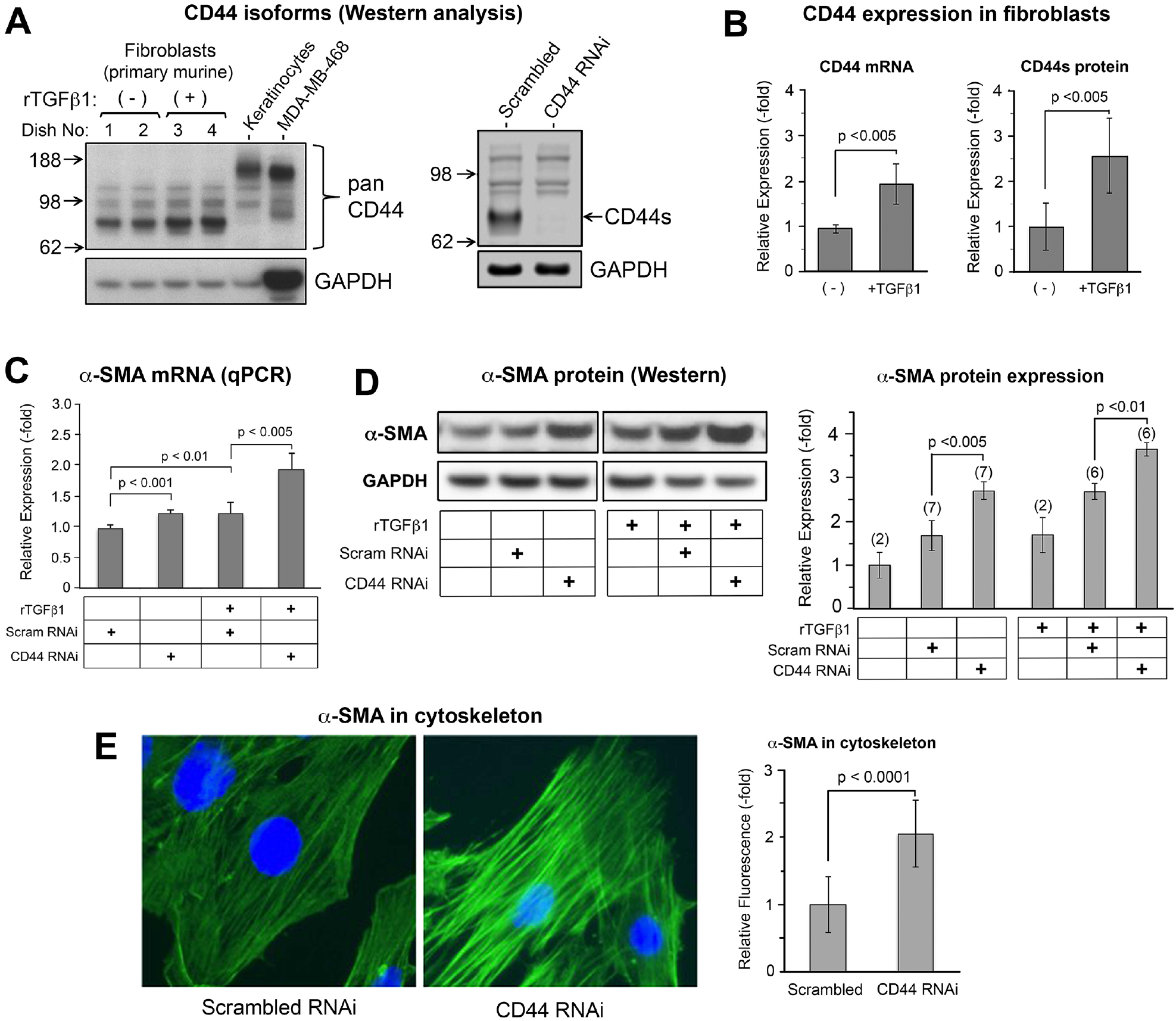
CD44s is the predominant isoform that controls α-SMA gene expression in primary murine skin fibroblasts. (**A**), Western blot of CD44 isoforms in fibroblasts, at baseline and 48 h after treatment ± 2 ng/mL of rTGF-β1 (*left panel*). Primary mouse keratinocytes and MDA-MB-468 breast cancer cells were positive controls for large CD44 isoforms (CD44E, and CD44v6, respectively). The band responsive to rTGF-β1 is CD44s. The other bands are nonspecific as shown by the knockdown experiment using CD44 RNAi (*right panel). GAPDH*, loading control. (**B**), Expression of CD44 mRNA by qPCR (*left panel*, n=5 cultures/bar) and CD44 protein by Western blot, (*right panel*, n=6 cultures/bar) in the absence or presence of rTGF-β1. (**C**), α-SMA mRNA levels by qPCR in cells transfected with control (*scram RNAi*) or CD44 siRNA (*CD44 RNAi*), ± 2 ng/mL of rTGF-β1 treatment for 48 h; mean ± SD of 4 independent experiments. (**D**), α-SMA protein levels in cells treated similarly to panel C; graph, mean ± SD of western blot experiments (number of pooled experiments shown above each bar). (**E**), α-SMA immunofluorescence in fibroblasts (*green*); nuclei counterstained with DAPI (*blue*). Graph, quantitation of α-SMA fluorescent staining intensity, mean ± SD, 10 images per condition pooled from 2 experiments.

To further clarify the role of CD44s in regulating α-SMA gene expression, we forcibly expressed a CD44s plasmid vector(37) in fibroblasts and observed downregulation of α-SMA expression on western blots (Figure 2A and 2B). Moreover, cotransfection of CD44s plasmid and CD44 RNAi effectively attenuated the increase in α-SMA levels induced by CD44 RNAi alone (Figure 2A and 2B), indicating that forced expression of the standard form of CD44 is inhibitory to α-SMA gene expression.

**Figure 2.**
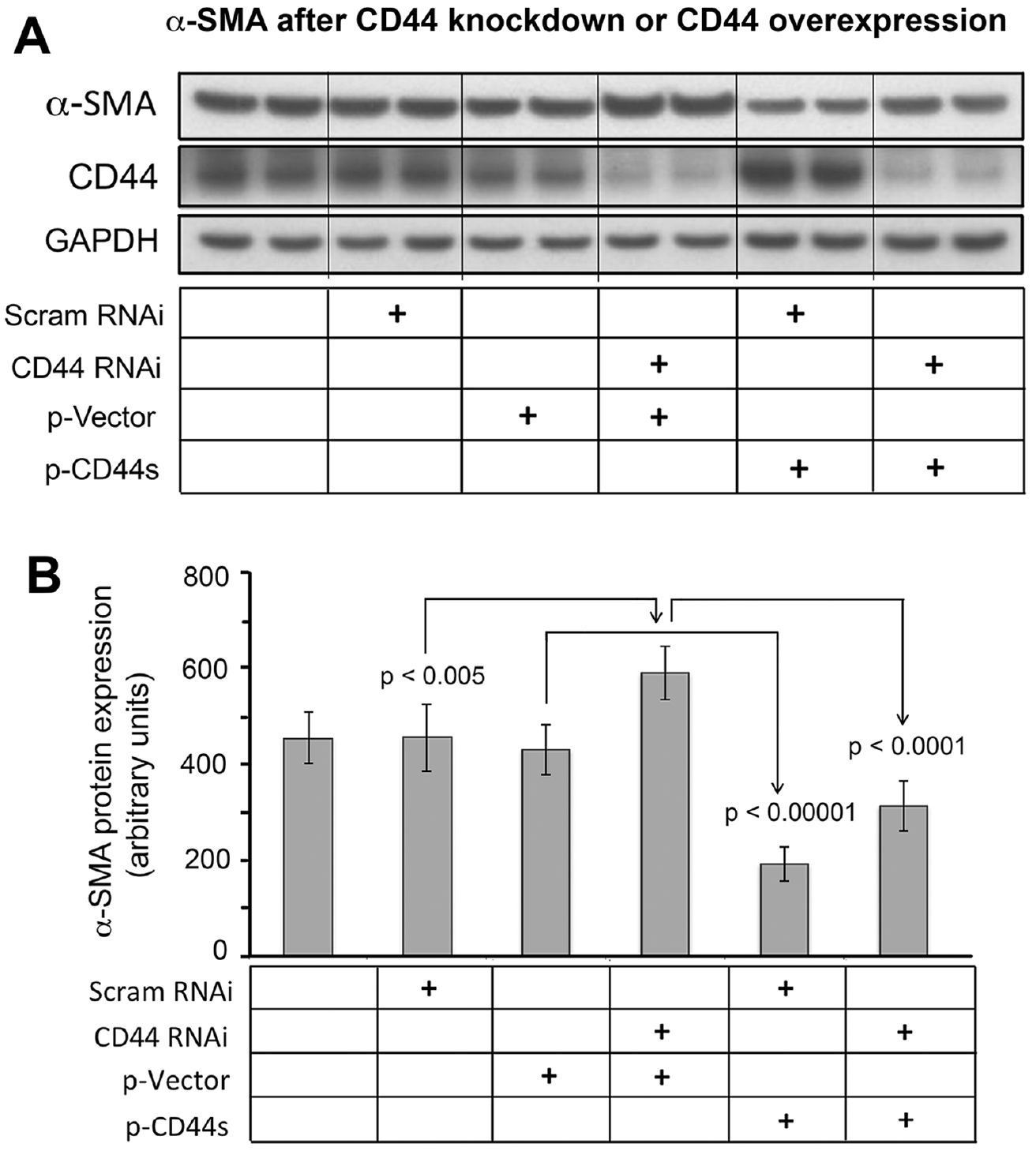
Effect of CD44 knockout or CD44 overexpression on α-SMA protein expression. *Scram RNAi*, scrambled siRNA. *CD44 RNAi*, transfection using CD44 RNAi. *p-Vector:* empty vector control. *p-CD44s*, forced expression using a CD44S plasmid. *Lower panel*, quantitative analysis of band intensities of α-SMA westerns; mean ± SD of 3 experiments.

### The upregulation of α-SMA gene expression induced by CD44 knockdown is not mediated by hyaluronan nor Has2

Since CD44 is the principal receptor for hyaluronan (HA) on plasma membranes of most cell types, including fibroblasts, and since HA is a critical regulator of pro-fibrotic signaling and MF differentiation in some tissues (see Introduction), we investigated whether HA is involved in regulating the expression of α-SMA in primary skin fibroblasts during manipulation of CD44 and/or TGFβ levels. Extracellular HA was removed by treating fibroblast cultures with hyaluronidase, and the efficient removal of extracellular HA was verified by immunofluorescent staining using HA-binding proteins as previously described (data not shown) (38). Cultures were subsequently treated with or without rTGF-β1 for 48 hours. Western analysis showed that α-SMA gene expression was not altered by hyaluronidase treatment, whether in the absence or presence of rTGF-β1 treatment (Figure 3A). Addition of 4-MU, a compound known to inhibit HA synthesis (38), also failed to alter α-SMA gene expression (Figure 3B). A third manipulation was to knock down Has2, the functionally predominant Has enzyme in murine skin fibroblasts (38). Knockdown efficacies of Has2 and CD44 were verified by q-PCR analyses (Figure 3C). An examination of HA levels by fluorophore-assisted carbohydrate electrophoresis (FACE) analysis (Figure 3D), showed that Has2 RNAi alone or together with CD44 RNAi significantly reduced the quantity of HA, both in the cell layer and in the conditioned media. However, despite marked reduction in HA levels due to Has2 knockdown, no effect on α-SMA expression was observed (Fig. 3E). CD44 knockdown on the other hand, either alone or in combination with rTGF-β1 treatment, led to consistent increases in α-SMA expression (Figure 3E), as previously observed.

**Figure 3.**
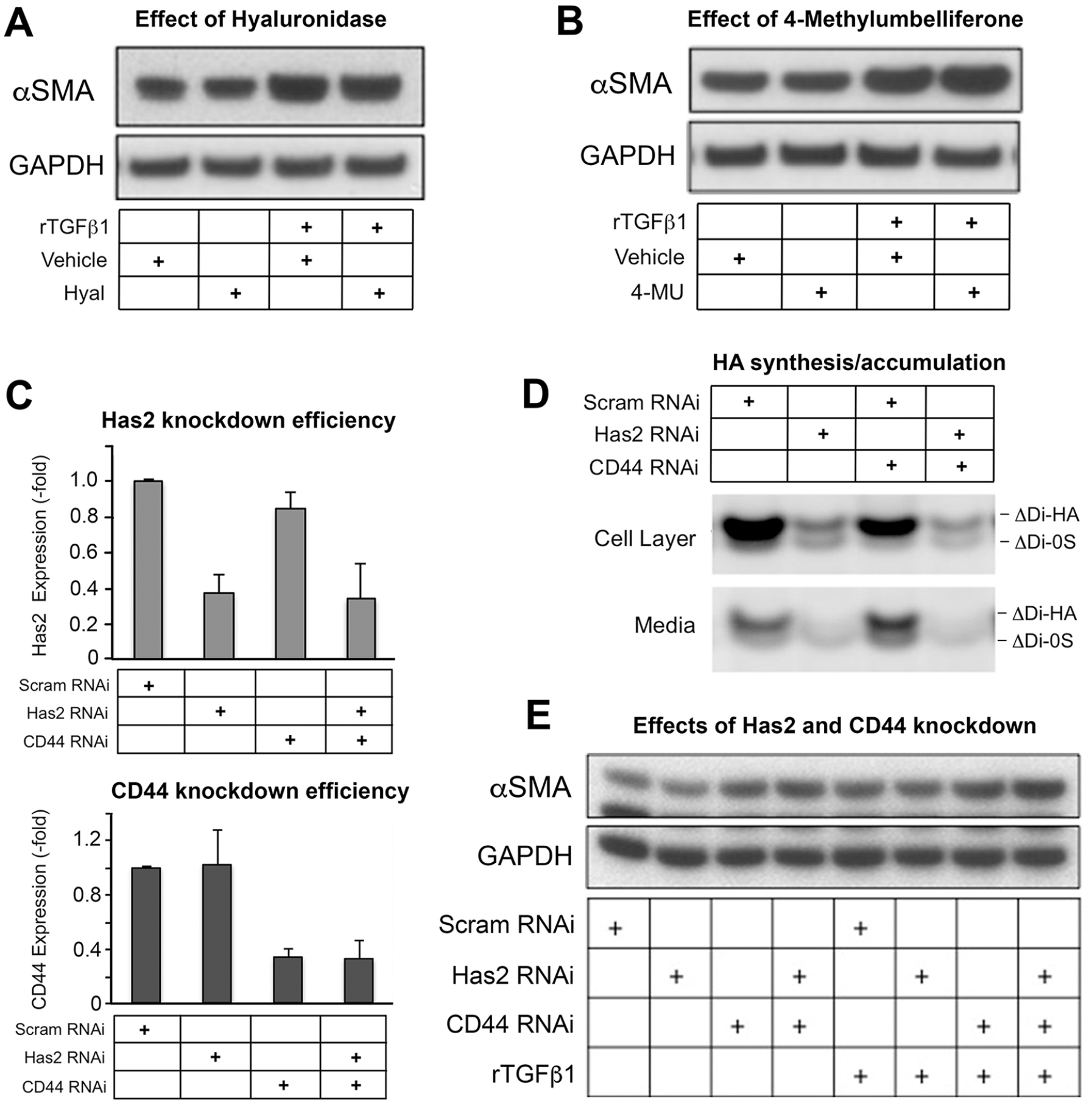
Inhibition of HA synthesis by 4-Methylumbelliferone (4-MU) or Has2 RNAi, or removal of extracellular HA with hyalruonidase (*Hyal*), has little effect upon α-SMA gene expression in murine skin fibroblasts. (**A**), SMA protein abundance of cells treated with *Hyal* followed by treatment with rTGF-β1, mean ± SD of 3 experiments. *Vehicle*, control. (**B**), α-SMA protein abundance in cells treated with 4-MU followed by rTGF-β1, mean ± SD of 3 experiments. *GAPDH*, loading control. (**C**), Knockdown efficiency with RNAi for Has2 or CD44 was validated by real-time qPCR; mean ± SD of 3 experiments). (**D**), A representative fluorescence-assisted carbohydrate electrophoresis (FACE) gel, showing HA disaccharide bands (Δ di-HA) from fibroblasts transfected with Has2 RNAi, and/or CD44 RNAi, or control (scram RNAi). Bands from the fibroblast cell layer and from the culture media are shown. (**E**), Western blot of α-SMA in fibroblasts transfected with Has2 RNAi and/or CD44 RNAi, followed by rTGF-β1 treatment; results are representative of 2 experiments.

### The regulatory effect of CD44 knockdown on α-SMA gene expression is dependent upon non-canonical TGF-β signaling that is mediated by p38MAPK

Given previous strong evidence for an important role for TGF-β signaling in regulating α-SMA gene expression, and for the involvement of CD44 (either positive or negative) upon TGF-βR–regulated signaling (see Introduction), we sought to determine which components of the TGF-β signaling cascade might be involved in the upregulation of α-SMA caused by CD44 knockdown. Fibroblasts were pre-treated with individual chemical inhibitors for membrane receptors (TGF-βR or EGFR), or with RNAi or chemical inhibitors for mid-level signaling molecules (SMADs, ERK, and p38), followed by transfection with CD44 RNAi and addition of rTGF-β1. As shown in Figure 4A, 4C, and 4D, inhibition of TGF-βR but not inhibition of EGFR suppressed α-SMA gene expression at baseline and abrogated the increased α-SMA gene expression induced by either CD44 RNAi or rTGF-β1 treatment.

**Figure 4.**
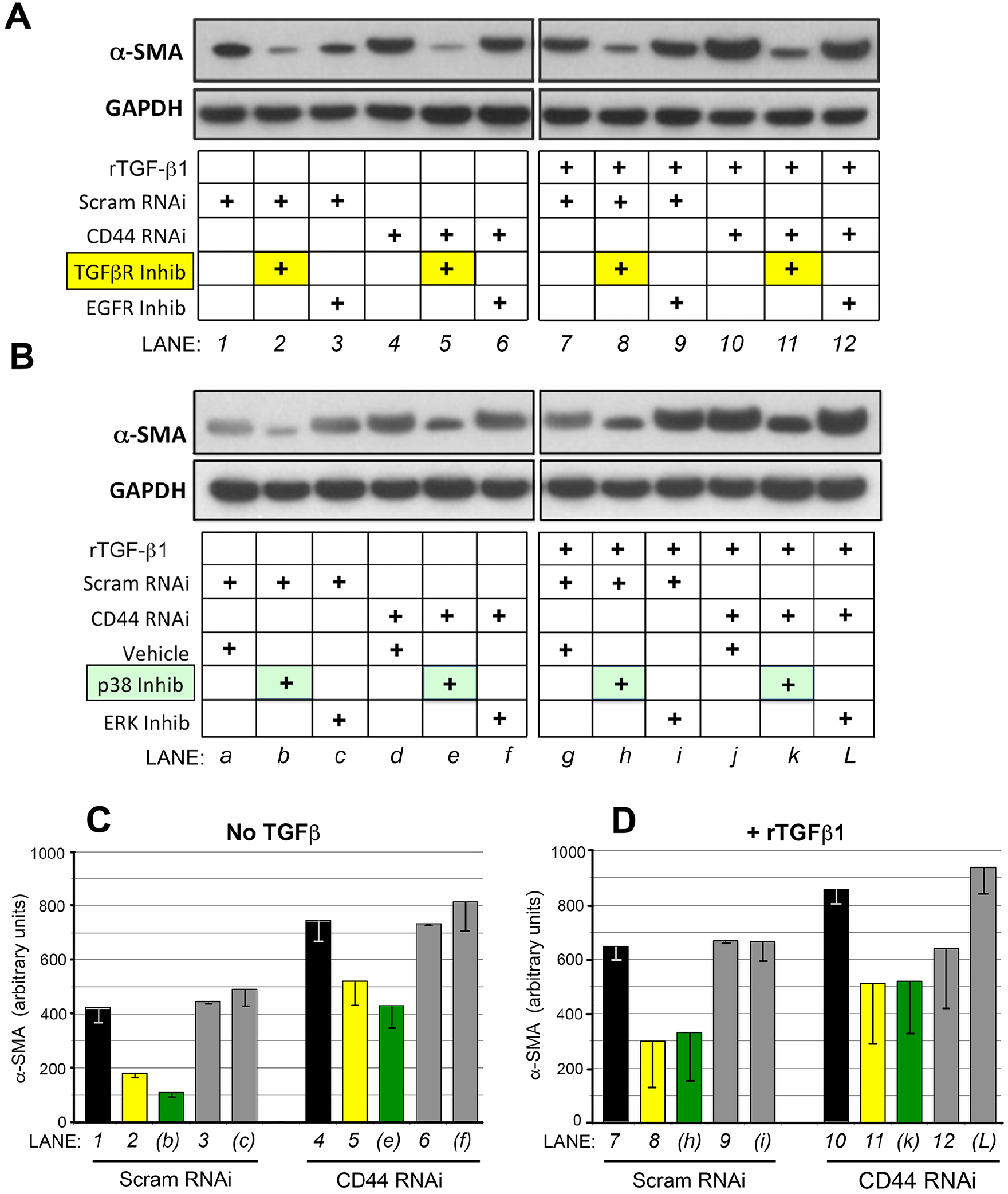
Chemical inhibitors reveal a role for TGF-β signaling in mediating the effects of CD44 RNAi upon α-SMA protein expression in fibroblasts. (**A**), Fibroblasts were pretreated for 4 h with TGF-β receptor inhibitor SB431542 (*<u>TGFβR Inhib</u>*), EGF receptor inhibitor AG1478 (*<u>EGFR Inhib</u>*), or vehicle only, then transfected with CD44 RNAi or scrambled RNAi (36 h incubation), followed by another 48 h in fresh media ± 2 ng/mL of rTGF-β1. Following these treatments, cells were harvested for western analysis of α-SMA and GAPDH (loading control). Results shown are representative of 4 experiments. (**B**), Fibroblasts were pretreated with p38MAPK inhibitor SB202190 (*<u>p38 inhib</u>*), ERK1/2 inhibitor U0126 (*<u>ERK inhib</u>*), or vehicle only for 4 h, then transfected with CD44 RNAi or scram RNAi for 36 h, followed by ± rTGF-β1 treatment for 48 h. Results representative of 4 experiments. (**C**) & (**D**), Densitometric quantification of α-SMA protein bands from western blots (± CD44 RNAi, ±TGF β1, and ± inhibitors) that were pooled from at least 2, but more typically from 3 or 4 experiments; mean ± SD.

To investigate how CD44 knockdown might affect SMAD-dependent TGF-β signaling, fibroblasts were transfected with either non-targeted or CD44 RNAi for 48 hours, then synchronized by serum starvation for another 24 hours, followed by addition of rTGF-β1 in the media. As shown in Supplementary Figure 1A, exposure to rTGF-β1 initiated a rapid increase in phosphorylated Smad2 and phosphorylated Smad3. However, CD44 RNAi had no effect on these phosphorylations relative to scrambled-RNAi controls, as analyzed by western blotting (Fig. S1A). Furthermore, Smad2 RNAi failed to alter α-SMA gene expression in murine skin fibroblasts either at baseline or in response to rTGF-β1 treatment, nor alter the induction caused by CD44 RNAi (Fig. S2), suggesting that SMAD-mediated TGF-β signaling is dispensable for the regulation of α-SMA expression in murine skin fibroblasts. CD44 RNAi also did not alter the overall pattern of rTGF-β1-induced phosphorylation of ERK1/2 (Fig. S1B). Knockdown of CD44 exerted a mild effect on p38, with a shift in the kinetics of rTGF-β1-induced phosphorylation (peak shifted from 30min in control cells to 6h in CD44 knocked down cells, with both returning to baseline by 12 hours) (Fig. S1C). Although CD44 did not show any major effect upon p38 phosphorylation, inhibition of p38 significantly suppressed α-SMA expression in the absence or presence of rTGF-β1, and effectively abrogated the increased α-SMA gene expression caused by CD44 knockdown (Fig. 4B, 4C and 4D), whereas inhibition of ERK1/2 had no such effect (Fig. 4B, 4C and 4D). This suggests that p38 activity is required for baseline α-SMA expression and also for upregulation of α-SMA expression when CD44 is knocked down. When combined, these results show no apparent role for EGFR nor SMADs, but instead show a clear involvement of TGFβR and p38 in the upregulation of α-SMA expression when CD44 is reduced in these cells.

### CD44 inhibition alters the ratio between G- and F-actins

Myofibroblast differentiation is accompanied by alteration of actin cytoskeleton architecture, including the incorporation of α-SMA into stress fibers (8). After observing that CD44 RNAi increases α-SMA gene expression in mouse skin fibroblasts, we wanted to know how the actin cytoskeleton might respond in cells receiving CD44 RNAi treatment. F-actin or G-actin was labeled and visualized with fluorescence-conjugated phalloidin or DNase I, respectively. As compared with control cells, either CD44 RNAi or rTGF-β1 treatment enhanced the formation of F-actin-positive stress fibers, and a combination of these two treatments yielded the best outcome (Figure 5A). On the other hand, these same treatments had the opposite effect on G-actin, causing a significant decrease in immunofluorescent staining intensity (Figure 5B). When G-actin was separated from F-actin by fractionation using an *in vivo* G-/F-actin assay kit, examination of both fractions by western analysis demonstrated a decrease in G-actin and an increase in F-actin in cells treated with either CD44 RNAi or rTGF-β1, relative to control cells that received nontargeted siRNA (Figure 5C). Consistent with the earlier findings using immunofluorescence (Figure 5A and 5B), the changes in Fig. 5C were most pronounced when CD44 RNAi and rTGFβ1 were administered together.

**Figure 5.**
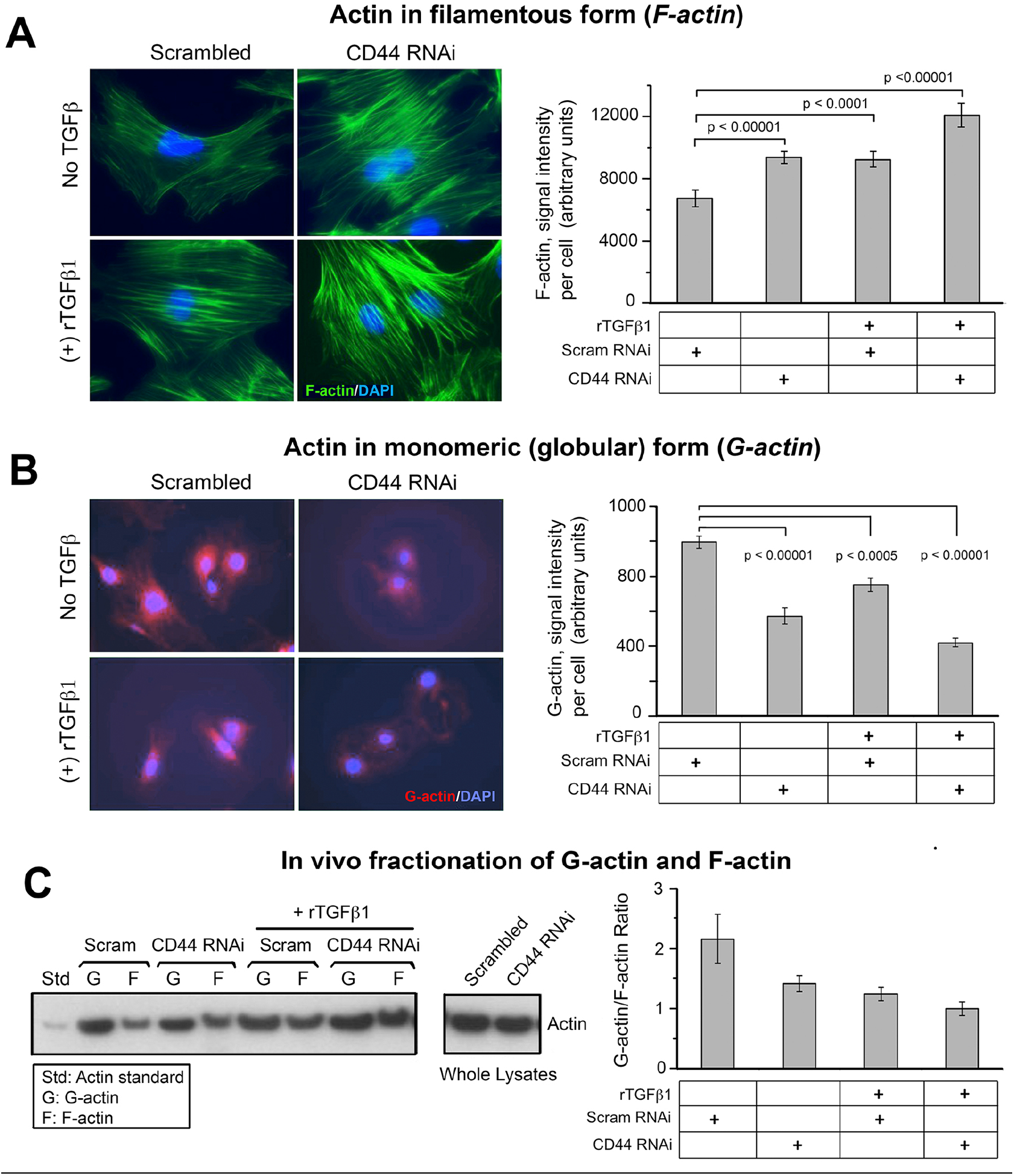
The effect of CD44 RNAi on the G-actin/F-actin ratio in fibroblasts. (**A**), images of F-actin stained with AF488-conjugated phalloidin, in cells transfected with CD44 RNAi or scrambled RNAi, ± 2 ng/mL of rTGF-β1 (48 h) (*left*). Quantitative analysis of the phalloidin signal using IPLab image analysis software; each bar, mean ± SD of ten high-power fields, 3 cells/field examined per condition, pooled from 2 experiments (*right*). (**B**), images of G-actin stained with AF594-conjugated DNAse I, in cells transfected as in Fig. 5A (*left*). Quantitative analysis of fluorescent intensity; each bar, mean ± SD of ten high-power fields/condition from 2 experiments (*right*). (**C**), representative western blot (*left*) and quantitative densitometric analysis (*right*) of G- and F-actin in fibroblasts transfected with scrambled or CD44 siRNA ± rTGF-β1 treatment for 48 h. F and G actin were isolated using the *in vivo* G-/F-actin assay (see Methods). Results are expressed as G-actin/F-actin ratio; each bar, mean ± half-range of 2 independent experiments.

### MRTF-A mediates the α-SMA induction effect of CD44 RNAi

It has been demonstrated by various groups that nuclear localization of MRTF and its binding to SRF play a crucial role in regulating the transcription of contractile genes including α-SMA (11,39,40). Among the three homologous members of the myocardin family (Myocardin, MRTF-A, MRTF-B), MRTF-A is expressed in nearly all adult tissues (41) and has a predominant role in regulating profibrotic signaling in skin tissue (42,43). We therefore focused on the role of MRTF-A (hereafter simply called “MRTF”). As shown in Figure 6A, expression levels of SRF and MRTF at both the protein and mRNA level were not affected by CD44 RNAi. However, CD44 knockdown did promote nuclear accumulation of MRTF as determined both by immunofluorescence assay (Figure 6B) and western blotting of MRTF in cytoplasmic and nuclear fractions (Figure 6D). SRF subcellular localization, on the other hand, was unaffected by CD44 RNAi (Figure 6C and 6D). Furthermore, CD44 knockdown resulted in increased binding between MRTF and SRF as determined by immunoprecipitation (Figure 6E). Lastly, knockdown of MRTF by RNAi (knockdown efficacy was verified by western analysis as shown in Fig. 7B) significantly suppressed the expression α-SMA in the absence or presence of rTGF-β1, and effectively abrogated CD44 RNAi’s ability to induce α-SMA gene expression (Figure 7A and 7C).

**Figure 6.**
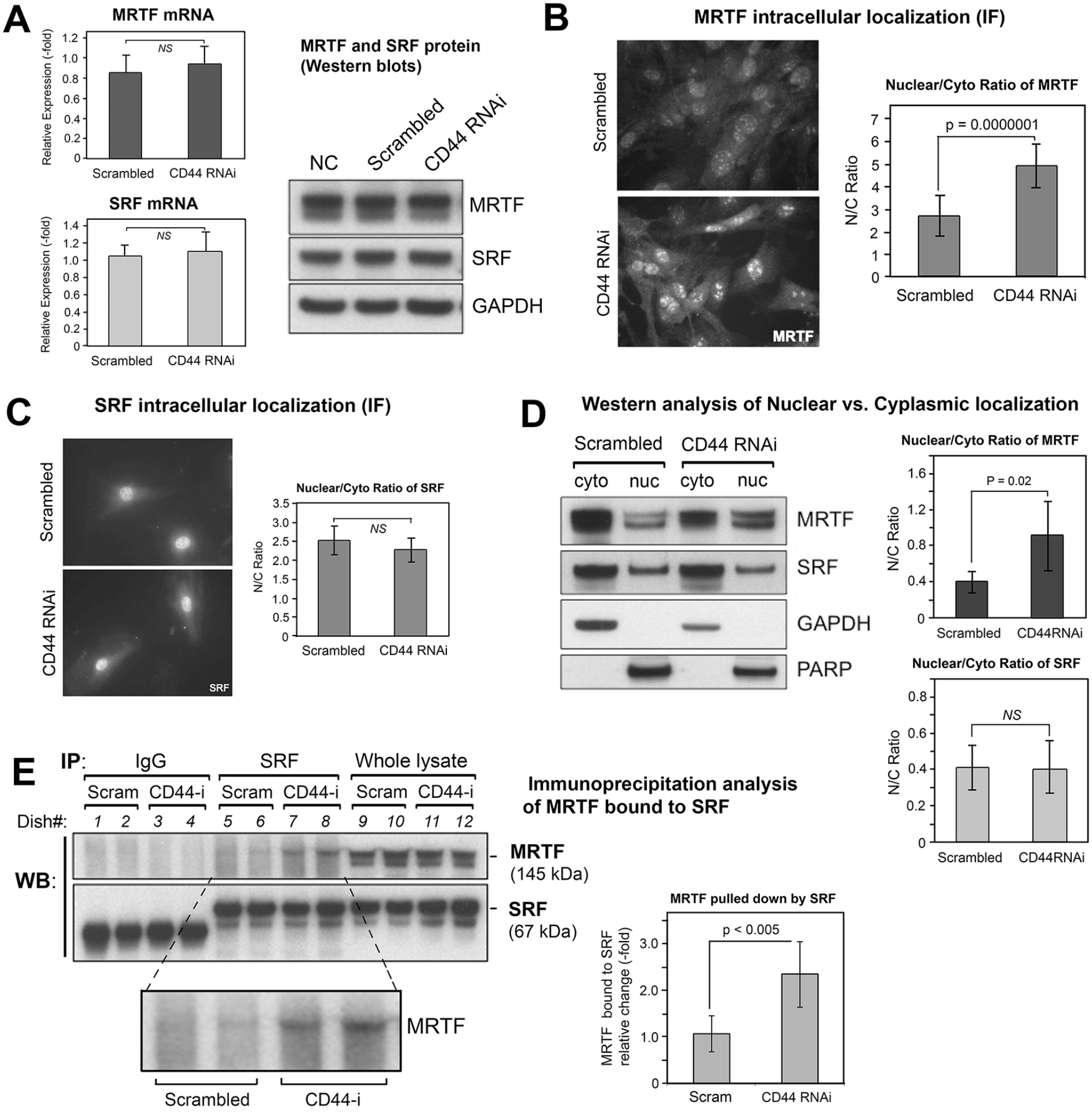
The effect of CD44 knockdown on gene expression and subcellular localization of MRTF and SRF. (**A**), mRNA levels of MRTF and of SRF by qPCR (*left*), mean ± SD of 3 experiments; protein levels of MRTF and SRF by western analysis (*right*), showing no difference in response to CD44 RNAi treatment. (**B**), MRTF-A immunofluorescent staining in fibroblasts transfected with CD44 RNAi or scram RNAi (*left*), and digital analysis of staining intensity to provide MRTF nuclear/cytoplasmic ratio (*right*); mean ± SD, 100 cells analyzed per condition. (**C**), SRF immunofluorescent staining in cells transfected as in **B** (*left*); digital analysis of SRF Nuc/Cyto ratio (*right*). (**D**), subcellular fractionation of MRTF in fibroblasts transfected with CD44 RNAi or scram RNAi. *Left*, representative western blot of MRTF from cytoplasmic (*cyto*) or nuclear (*nuc*) fractions, along with GAPDH and PARP as controls for purity of the *cyto* and *nuc* fractions, respectively. *Right*, Nuc/Cyto ratio of MRTF, determined from densitometric scanning (IPLab) of bands in two independent experiments, mean ± S.D. (**E**), immunoprecipitation analyses to assess binding between SRF and MRTF in fibroblasts transfected with CD44 or scram RNAi. An anti-SRF antibody (*SRF*) or normal rabbit immunoglobulin (*IgG*) was added to the cell lysates to pull down protein complexes, and the membrane probed with anti-MRTF or anti-SRF antibodies (*left*). Quantitative analysis of the SRF-bound MRTF band (*inset*) is shown in the graph (*right*); bars, mean ± SD of two experiments with duplicate samples.

**Figure 7.**
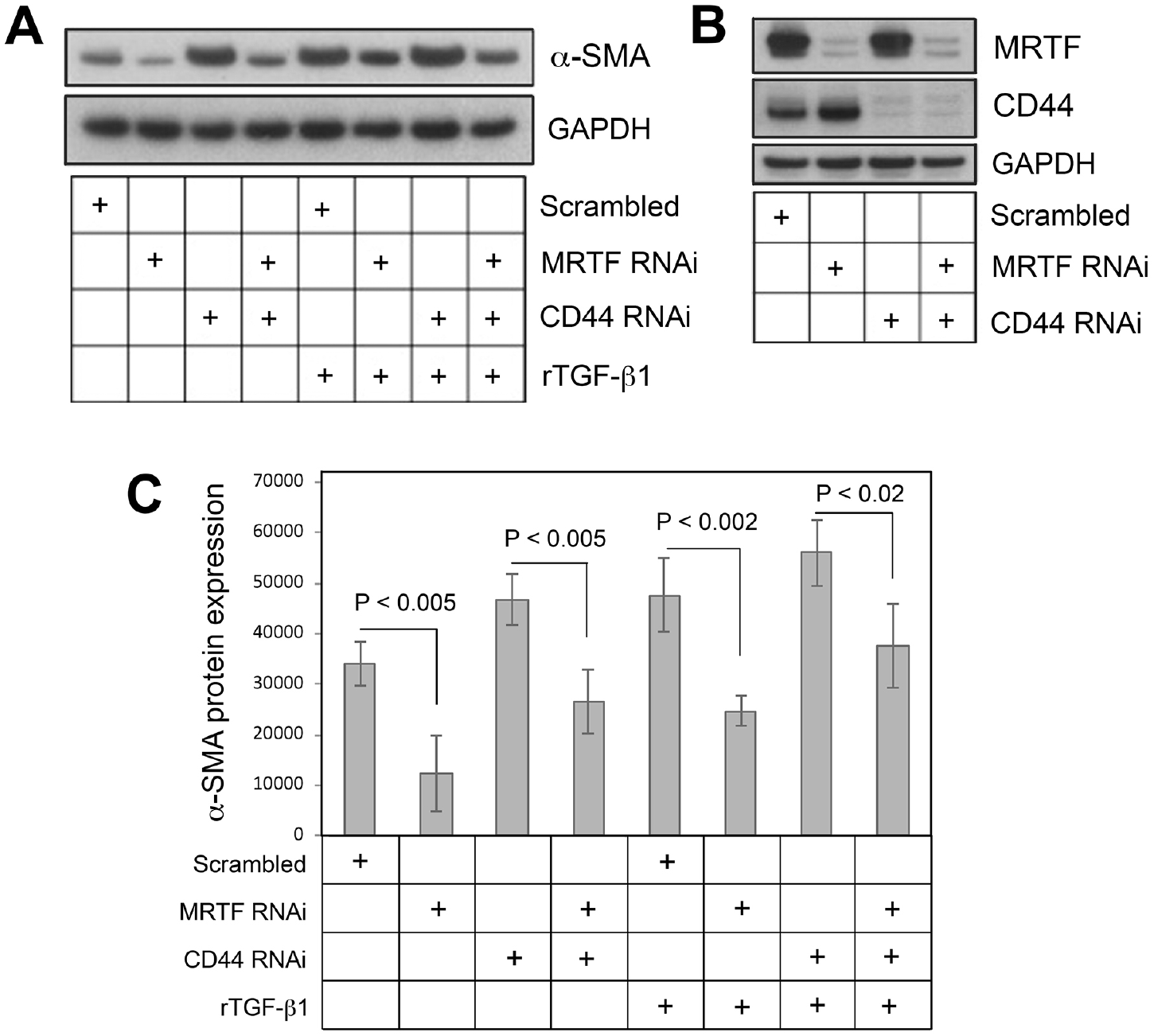
Knockdown of MRTF partially abrogates the α-SMA upregulation induced by CD44 knockdown or rTGF-β1 treatment. (**A**), α-SMA western blot in fibroblasts transfected with MRTF RNAi and/or CD44 RNAi, ± rTGF-β1. (**B**), knockdown efficacy (western blot) of MRTF and CD44 for the experiment shown in panel A. (**C**) Densitometric analyses of α-SMA western blots using IPLab imaging software, normalized to GAPDH, pooled from 4 experiments; bars, mean ± range.

### The regulatory effect of CD44 RNAi and TGF-β receptor signaling on the subcellular localization of MRTF is dependent on p38MAPK activity

Nuclear accumulation of MRTF and binding to SRF play a central role in regulating α-SMA gene expression. As shown in Figure 8A and 8B, either CD44 RNAi or rTGF-β1 treatment was able to induce nuclear accumulation of MRTF in primary murine dermal fibroblasts. Either effect could be prevented by pretreatment of the cells with SB202190 (p38-i), a chemical inhibitor of p38MAPK. The expression level of MRTF was not altered by treatment with SB202190 as determined by Western analysis (data not shown). During p38MAPK inhibition, MRTF appeared completely absent from a majority of nuclei. These results indicate that the regulatory effects of both CD44 RNAi and TGF-β1 upon subcellular localization of MRTF are dependent on p38MAPK activity.

**Figure 8.**
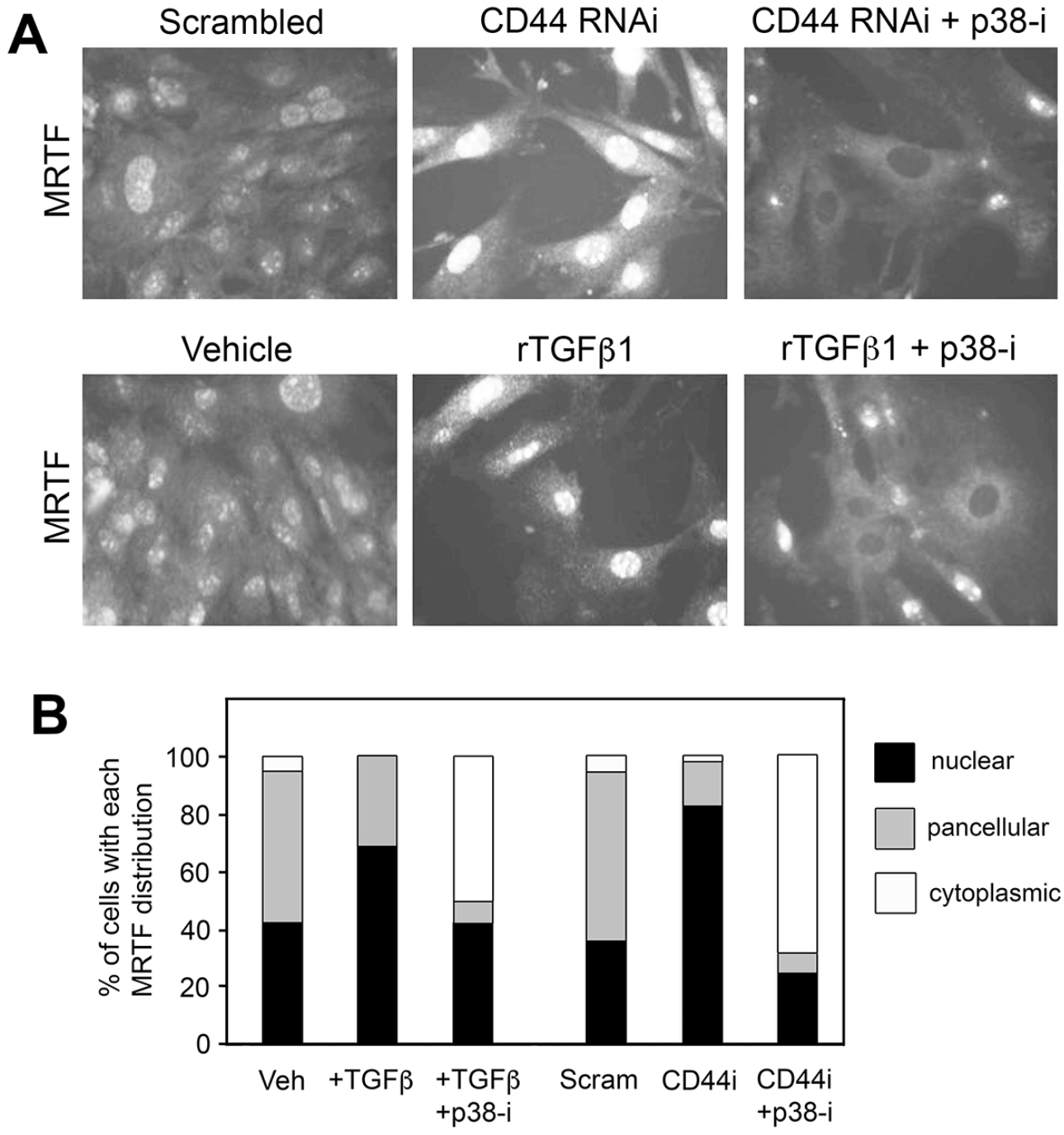
Effects of CD44 RNAi and TGF-β receptor signaling on the subcellular localization of MRTF-A are dependent upon the p38MAPK activity. (**A**), Upper panel: immunofluorescently stained MRTF in fibroblasts treated with either Scrambled siRNA, or CD44 RNAi in the presence or absence of SB202190, a p38MAPK inhibitor (*p38-i*). Lower panel: immunofluorescently stained MRTF in fibroblasts treated with either vehicle control, or rTGF-β1 in the presence or absence of SB202190 (*p38-i*). (**B**), Analysis of subcellular MRTF, in which each cell was scored for its predominant intracellular distribution (cytoplasmic, nuclear, or pancellular). Each bar represents 100 to 300 cells analyzed in 6 images per treatment condition.

## DISCUSSION

The significance of the study presented here is that to our knowledge, this report is the first description of a direct regulatory effect of CD44 on α-SMA gene expression via an actin/MRTF/SRF-dependent pathway. Our working model for α-SMA regulation in mouse fibroblasts is summarized in Figure 9. Most previous studies that investigated CD44’s role in pro-fibrotic signaling and regulation of α-SMA expression focused on characterizing the interactions between CD44 and EGFR, or between CD44 and TGFβR in the plasma membrane, but contradictory observations were generated depending upon the cell type and experimental conditions (see Introduction). Thus, CD44 was stimulatory for α-SMA expression in some systems and inhibitory in others. Curious about this discrepancy, we examined primary murine skin fibroblasts and found that knockdown of CD44 actually promoted α-SMA expression in these cells (notably similar to what Porsh *et al* had reported in human dermal fibroblasts) (32). We then sought to determine whether this effect is mediated by EGFR or TGFβR signaling and found that knockdown of CD44 had little to no impact upon either pathway, at least in terms of canonical signaling. Blockade of EGFR or ERK1/2 did nothing to blunt the effect of CD44 knockdown, and ERK1/2 phosphorylation was unaffected by CD44 RNAi, suggesting no involvement of EGFR signaling. Regarding TGFβR signaling, we found that CD44 knockdown did not alter the phosphorylation of Smad2/3, and had only mild effects on phosphorylation of p38MAPK, suggesting that CD44 RNAi does not directly modulate TGFβR activation. The mild effect on p38MAPK might be achieved via other pathways or could be an indirect effect. Our experimental approach of knocking down Smad2 failed to rescue the increase in α-SMA expression caused by CD44 RNAi, suggesting that the classical TGFβR/Smad pathway is dispensable in regulating α-SMA expression in primary murine skin fibroblasts. These findings motivated us to explore other pathways that might be responsible for mediating CD44’s regulatory effect on α-SMA. We observed that increased α-SMA gene expression after CD44 depletion was accompanied by a pronounced shift in the actin cytoskeleton, with a significant increase in F-actin filaments and a decrease in monomeric G-actin in the cytosol, accompanied by increased nuclear accumulation of MRTF and enhanced MRTF-SRF binding. Moreover, knockdown of MRTF markedly reversed the up-regulated gene expression of α-SMA induced by CD44 RNAi, suggesting that MRTF mediates the regulatory effect of CD44 RNAi on α-SMA.

**Figure 9.**
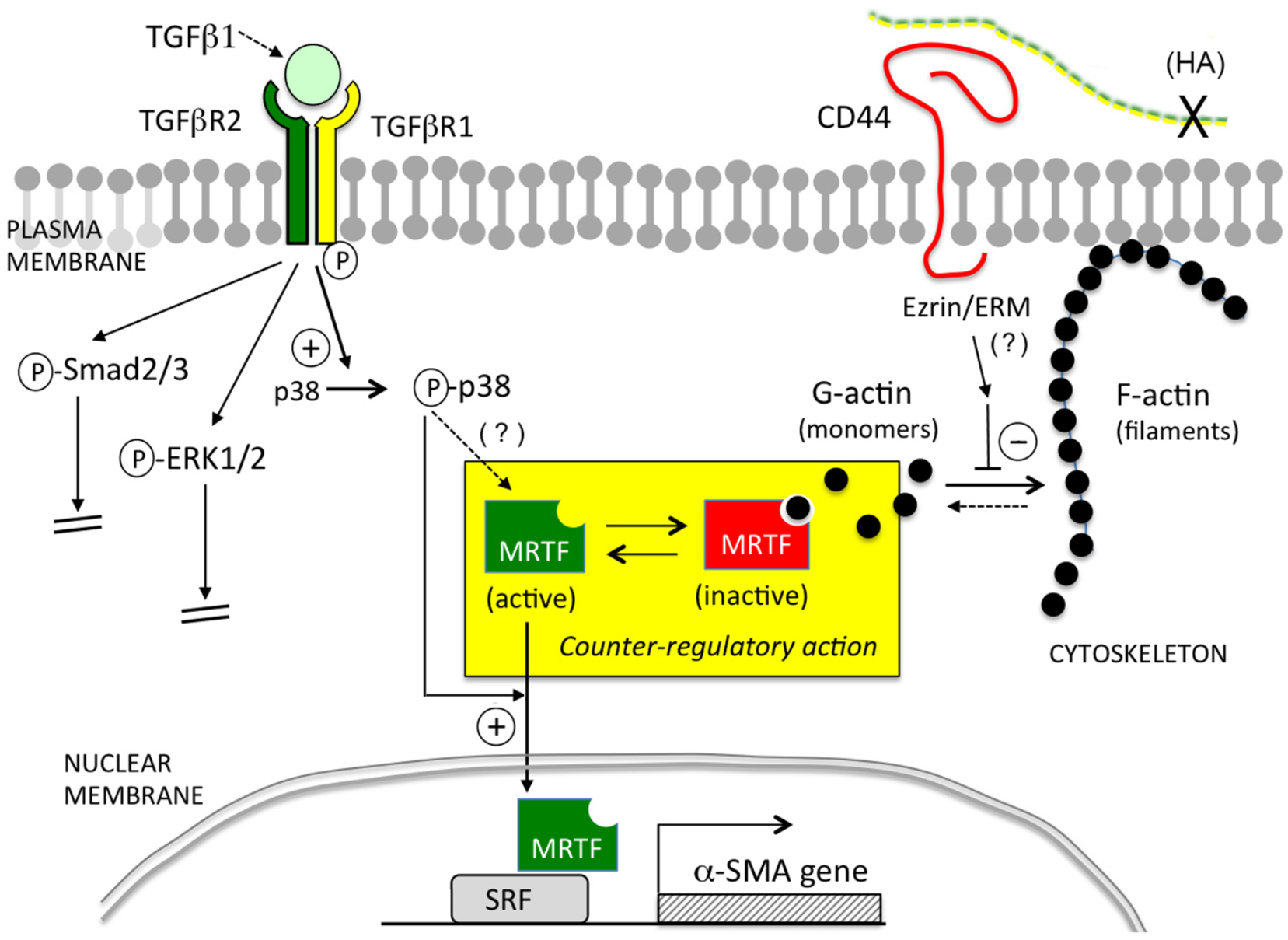
Summary Diagram. Cartoon illustration of the pathways examined in this study. In the context of primary murine dermal fibroblasts, TGF-β/TGFβR-mediated signaling positively regulates α-SMA expression via p38MAPK instead of via Smad2/3 or ERK1/2. CD44 negatively regulates α-SMA expression by inhibiting actin polymerization, leading to more globular G-actin in the cytoplasm which binds and sequesters MRTF, preventing MRTF from entering the nucleus and binding to SRF. Activation of p38MAPK is required for upregulation of α-SMA expression caused by either TGFβR signaling or knockdown of CD44.

The fact that MRTF (together with its obligatory binding partner, SRF) can control myofibroblast differentiation was previously documented in cells from various tissues including cardiac, lung and skin fibroblasts, and kidney and ovarian epithelial cells (44). Crider et al showed that knockdown of MRTF-A/B effectively reduced the expression of smooth muscle-specific proteins (α-SMA, SM22-α, etc.) induced by TGF-β1 treatment, whereas forced expression of a constitutively-active MRTF-A construct markedly increased the expression of α-SMA and SM22-α, along with contractile function in rat embryonic fibroblasts (39). In skin, Velasquez et al showed that treating mice with a small molecule, Isoxazole, promoted wound healing *in vivo* by regulating MRTF stability and activity (40). Our results showed that MRTF is crucial not only for α-SMA expression in primary murine dermal fibroblasts but also for mediating CD44’s regulatory effect. This effect of CD44 upon the functionality of MRTF appears to be mediated through the actin cytoskeleton system, as shown by our data that demonstrate shifts in the G-to-F actin profile (Fig.4).

The nature of the link between MRTF, actin, and α-SMA transcription is well-established from previous studies. MRTFs contain a basic region (B1) and an adjacent Glu-rich domain through which they associate with SRF (10,45), and an N-terminal domain (NTD) that contains three Arg-Pro-X-X-X-Glu-Leu (RPEL) motifs that mediate MRTF binding with G-actin. At baseline, when levels of actin polymerization are low, monomeric G-actins bind to MRTF via the RPEL motifs and sequester MRTF in the cytoplasm. Upon stimulation by growth factors and cytokines, activation of Rho GTPases (RhoA, Rac, and Cdc42) induces actin polymerization through conversion of G-actin into F-actin filaments, thus freeing MRTF to enter nuclei and bind to SRF(11). In addition to this well-established regulatory mechanism, phosphorylation on certain residues of MRTF regulates MRTF subcellular localization and function. Both ERK and p38MAPK can phosphorylate MRTF at certain sites. ERK-mediated phosphorylation of MRTF-A at serine 454 inhibits nuclear localization of MRTF induced by serum (46), and p38MAPK/MK2 can stimulate MRTF-A phosphorylation at serine 312 and serine 333 (47). MRTF phosphorylation by p38MAPK in HeLa cells or MEF cells (46,47) did not appear to affect MRTF binding to actin nor its subcellular localization, but such effects could be cell-context dependent since regulation of MRTF nuclear accumulation and α-SMA promoter activity via activation of p38MAPK was demonstrable in porcine kidney epithelial cells (48). Here in our study, chemical inhibition of p38MAPK blocked α-SMA expression at baseline, and aborted the α-SMA induction effect caused by CD44 RNAi. Although CD44 knockdown did not appear to affect p38MAPK phosphorylation status, chemical inhibition of p38MAPK led to marked cytoplasmic sequestration of MRTF, effectively abrogating the increased α-SMA gene expression induced by CD44 RNAi. This indicates that p38MAPK kinase activity is required and permissive for CD44’s regulatory effect on α-SMA gene expression via MRTF. Our observation is in harmony with findings from many other groups that show p38MAPK plays a crucial role in mediating TGF-β-induced MF differentiation and tissue fibrosis(16-18).

We showed in this report that in primary murine fibroblasts, only the shortest CD44 isoform known as CD44 standard (CD44s) is detectable by Western analysis, as opposed to other variants that may be produced by alternative splicing in certain tissues, such as variant isoforms v6 or v7/8 that play a role in regulating cellular behavior in certain fibroblasts (49,50). We also demonstrated that forced expression of CD44s suppressed α-SMA gene expression at baseline and abrogated the increased α-SMA gene expression caused by CD44 RNAi, further strengthening our argument that CD44s plays a predominant role in regulating α-SMA gene expression in primary murine skin fibroblasts. How CD44 is regulating the G-to-F actin cytoskeletal changes that regulate MRTF, and why such a relationship was not widely recognized before, are very interesting questions. One reason may be that CD44 manipulation (e.g., CD44 deletion or overexpression) typically yields only mild phenotypes. For example, mice with global CD44 deletion showed only minor abnormalities in leukocyte behavior (19), although injury brought out more significant differences, such as a delay in wound reepithelialization in mice with epidermal-specific deletion of CD44 (CD44 floxed-k14Cre) (51). In wounded BALB/c mice, CD44 deficiency did not alter skin wound closure rates, but did attenuate FAP-positive fibroblast accumulation during wound healing along with increased fibrillar collagen accumulation (52).

An important unanswered question is how CD44 might be controlling the G-to-F actin cytoskeletal filament changes that regulate MRTF. A strong connection between CD44 and regulation of cytoskeletal dynamics has been demonstrated in many studies. For example, CD44 has a well-established role as an organizer of the cortical actin cytoskeleton through direct interactions with ERM (Ezrin/Radixin/Moesin)(19,53) or other membrane-associated cytoskeletal proteins such as ankyrin(54). In addition, CD44 may also regulate actin cytoskeleton by modulating the activity of Rho GTPases (55-63). For example, CD44 can mediate cell adhesion (64-66). Disruption of epithelial cellcell contacts by calcium deprivation can trigger the activation of Small GTPases and lead to nuclear accumulation of MRTF in a Rho-, or Rac1- or p38-dependent manner(48,64-68). In our system, CD44 knockdown might disrupt cell-cell contacts, triggering GTPase activation and increasing nuclear MRTF. Protein Kinase C (PKC), a family of serine/threonine kinases, are another set of candidates for involvement since they mediate intracellular signaling, remodel cytoskeletal microfilaments by interacting with various actin-binding proteins (69), and can regulate the expression and function of CD44 (70-73). In future follow-up studies, any of the above molecules might be considered as candidates for understanding how CD44 inhibition (or overexpression) regulates changes in the G-/F-actin ratio and subcellular relocalization of MRTF.

Another surprising finding was the absence of any apparent role for HA in the effects of CD44 in our system. Studies from researchers at Cardiff University established both HA and CD44 as important facilitators of TGF-β-driven MF differentiation in human lung fibroblasts (see Introduction) which was mediated by the canonical Smad2/3-dependent pathway (24,25). Conversely, Evanko et al showed that reduction or removal of HA led to enhanced MF differentiation in human lung fibroblasts (along with increased fibronectin and type I collagen deposition) suggesting that HA has a negative effect on α-SMA gene expression and MF differentiation (74). In our study we found that CD44 inhibits MF differentiation in primary murine fibroblasts, independently of HA. This is not unprecedented. For example, Schmitt and colleagues showed that in human embryonic kidney cells, CD44 is a positive regulator of canonical Wnt signaling by regulating the phosphorylation and membrane localization of low-density lipoprotein receptor-related protein 6 (LRP6), and this regulation was not affected by either hyaluronidase treatment or by antibodies that block HA/CD44 binding, suggesting that CD44-mediated amplification of Wnt signaling is independent of HA. The reason for these discrepancies remains to be elucidated, but it may be that differences in species, type of tissue, and cell culture conditions are important contributing factors. Also, it is well recognized that HA is not the only ligand for CD44. CD44 can bind to osteopontin, fibronectin, type I collagen, fibrin, P-selectin, and E-selectin, any of which might theoretically play a role in HA-independent CD44-mediated signaling (19,20,75).

In conclusion, the present study shows that CD44 regulates α-SMA gene expression in primary murine dermal fibroblasts by controlling actin cytoskeleton dynamics and subcellular localization of MRTF in an HA-independent manner. The subcellular localization of MRTF is also susceptible to p38MAPK activity, which is a non-canonical downstream target of TGFβR signaling. These two pathways intersect at the level of MRTF and mediate the regulatory effect of CD44 on α-SMA gene expression. These findings add new components to the complexity of how CD44 interacts with membrane receptor kinases such as TGFβR, and provide novel insights about how CD44 may regulate pro-fibrotic signaling and myofibroblast differentiation. Importantly, further experiments are needed to elucidate which ligand(s) bind to CD44 to allow regulation of α-SMA expression, and which pathway(s) or molecule(s) are involved in mediating CD44’s role in the dynamic changes in MRTF and the actin cytoskeleton in primary mouse skin fibroblasts.

## EXPERIMENTAL PROCEDURES

### Primary Cell Culture

Primary mouse dermal fibroblasts were isolated from the skin of 2 to 3 day old pups from wild-type C57BL/6J mice (JAX Laboratories, Bar Harbor, ME) following a previously published protocol, as approved by the hospital’s animal review committee (Cleveland Clinic IACUC) (38). Briefly, the entire trunk skin was removed and incubated overnight in 0.25% trypsin without EDTA, followed by mechanical separation of epidermis from dermis. To isolate fibroblasts, the dermis was finely diced and incubated with 400 U/ml of collagenase type I (Worthington Biochemical, Lakewood, NJ, USA) for 30 minutes at 37 °C, then with 100 U/ml of DNAase I (Worthington Biochemical, Lakewood, NJ, USA) for 10 minutes at 37 °C. The suspension was then passed through a 100μM cell strainer and centrifuged at 300 rpm for 3 min. The fibroblasts were collected and cultured in Dulbecco’s Modified Eagle’s medium (DMEM) containing 10% fetal bovine serum (FBS), 1% penicillin/streptomycin, and 1.0 g/L of glucose. The cells were maintained at 37°C with 5% CO2. For all experiments, subconfluent cells from passage 2 or 3 were used.

### Inhibition of HAS2, CD44, Smad2, and MRTF gene expression by RNA interference (RNAi)

ON-TARGETplus SMARTpool Small interfering RNA (siRNA) and control siRNA was purchased from Dharmacon (Lafayette, Colorado): CD44 [*L-041132-01*], Has2 [*L-042589-01*], Smad2 [*L-040707-00*], MRTF/Mkl1 [*L-054350-00*], and nontargeted scrambled siRNA [*D-001810-10*]. The siRNAs were reverse transfected into mouse skin fibroblasts using Lipofectamine RNAiMAX (Invitrogen/Thermo Scientific, Carlsbad, CA) as previously described (38). Briefly, for a 6-well tissue culture plate format, 30 pmol of siRNA was mixed with 8μL of Lipofectamine RNAiMAX in 500 ml of Opti-MEM with a final duplex concentration of 100nM. The siRNA/transfection reagent mixture was added to the plate first, followed by seeding the fibroblasts at a density of 2.2 × 10^5^ per well. The cells were re-fed with fresh antibiotic-free medium supplemented with 10% FBS 24 h after transfection and further incubated for another 36~48 hours before harvesting for assays. The knockdown efficacy of target gene expression by RNAi was confirmed using quantitative real-time PCR and western blotting of the targeted gene product.

### CD44 plasmid purification and transfection

The plasmid for the standard form of CD44 (CD44s) was a generous gift from Dr. Veronique Orian-Rousseau(37). The CD44s insert in the plasmid was confirmed by DNA sequencing. The CD44s plasmid or an empty pcDNA3.1 mammalian expression vector (Invitrogen/ThermoFisher Scientific, Carlsbad, CA) was amplified using One Shot^™^ Top10 chemically competent E. Coli (Cat# C4040-10, Invitrogen/Thermo Scientific, Carlsbad, CA) and purified using the QIAGEN Plasmid Maxi Kit (Cat# 12162, QIAGEN, Hilden, Germany), then transfected into cells using Lipofectamine 3000 (Invitrogen/Thermo Scientific, Carlsbad, CA), according to the manufacturers’ instructions.

### RNA isolation and quantitative Real-time RT-PCR

RNA was prepared from fibroblasts using Trizol reagent (Life Technologies, Carlsbad, CA) per the manufacturer’s instruction. cDNA was then synthesized using random primers and Superscript III reverse transcriptase (Life Technologies, Carlsbad, CA). For real-time PCR, TaqMan gene expression probes for Has2 (Cat# Mm00515089_m1), CD44 (Cat# Mm01277161_m1), α-SMA (Mm01546133_m1), MRTF-A (Mm00461840_m1), SRF (Mm00491032_m1), and 18S ribosomal RNA (endogenous control, Cat# 4333760F) were purchased from Applied Biosystems/Life Technologies (Foster City, CA). The mRNA transcript levels of all target genes were measured in triplicate on an Applied Biosystems 7500 Real-time PCR system, and calculated using the 2-ΔΔCt method. The mRNA levels were presented as -fold differences relative to untreated normal controls.

### In vivo G-/F-actin assay

Soluble G-actin was separated from filamentous F-actin using the *In Vivo G-actin/F-actin Assay Kit* (Cat# BK037, Cytoskeleton, Inc., Denver, CO) according to the manufacturer’s instructions. Briefly, fibroblasts were harvested by scraping into lysis buffer supplemented with ATP and centrifuged at 350 × g to pellet cell debris. The supernatant was transferred to a new tube and centrifuged at 100,000 × g at 37 °C for 1 h to pellet the F-actin. After the G actin-containing supernatant was transferred to a new tube, a depolymerization buffer was added to the pellet and incubated on ice for 1 h to depolymerize the F-actin. Following depolymerization, a 5X SDS sample buffer was added to the G-actin and F-actin-containing samples and heated at 95 °C for 5 min to dissolve the proteins, which were then analyzed by western blotting.

### Subcellular fractionation

Nuclear and cytosolic fractions were separated using the *Nuclear/Cytosol Fraction Kit* from BioVision (Cat# K266-25; Milpitas, CA) per manufacturer’s instructions. Briefly, cultured fibroblasts were harvested and dissolved in cytosol extraction buffer A, vortexed vigorously for 15 s, and ice-cold cytosol extraction buffer B added to each tube and vortexed for an additional 5 s. The suspension was centrifuged for 5 min at 16000x g, and the supernatant (cytosolic extract) transferred immediately to a new tube. The pellet (nuclei) was resuspended in ice-cold Nuclear Extraction Buffer Mix and vortexed for 15 s every 10 min for a total of 40 min, followed by centrifugation at 16,000 × g for 10 min. The supernatant (nuclear extract) was transferred immediately to a new tube.

### Immunoprecipitation

Fibroblasts were scraped off the culture dishes and lysed in RIPA (Radioimmunoprecipitation assay) buffer (25mM Tris, 150mM NaCl, 0.1% sodium dodecyl sulfate, 0.5% sodium deoxycholate, 1% Nonidet P-40, 0.5mM EDTA) supplemented with protease inhibitor cocktail (EMD/Millipore, Gibbstown). A 10 μL aliquot of either anti-SRF antibody (Catalog # 5147, Cell Signaling Technology, Danvers, MA) or IgG control (Cat# 2729, Cell Signaling Technology) was added to 500 μg/500 μL of lysate and incubated at 4 °C overnight. After the incubation, 20 μL of pre-equilibrated protein A/G agarose beads (#SC2003, Santa Cruz Biotechnology, Dallas, TX) was added to each sample and incubated with rotation at 4 °C for 2 h, followed by washing with RIPA buffer three times. Protein elution was done by heating the beads in 45 μL of 5x Laemmli sample buffer at 95 °C for 5 min, followed by centrifugation at 15,000 rpm for 15 s. The supernatant containing the eluted proteins was transferred to a new tube. Eluted proteins were probed for SRF and MRTF using western blotting (see below).

### Western Blot Analysis

Primary antibodies purchased from Cell Signaling Technology were anti-phospho-Smad2 (Ser465/467, Cat# 3108), anti-Smad2 (Cat# 5339), anti-phospho-Smad3 (Ser423/425, Cat# 9520), anti-Smad3 (Cat# 9523), anti-phospho-ERK1/2 (Cat# 9102), anti-phospho-p38MAPK (Thr180/Tyr182, Cat#9212), anti-p38MAPK (Cat# 9212), anti-MRTF/MKL1 (Cat# 14760), anti-SRF (Cat# 5147), and anti-PARP (Cat# 9542). Primary antibodies purchased from Santa Cruz Biotechnology were anti-CD44 (Cat# sc-7051-R) and anti-GAPDH (Cat# sc-25778). Polyclonal rabbit anti-α-smooth muscle actin (Cat# ab5694) was purchased from Abcam (Cambridge, MA). Secondary antibodies including goat anti-rabbit or goat anti-mouse IgG conjugated with horseradish peroxidase (HRP) were obtained from Jackson ImmunoResearch Laboratories (West Grove, PA). At the end of treatments, cells were scraped off the culture plates and lysed in RIPA buffer supplemented with protease inhibitor cocktails (Millipore) and were then diluted in 4X NuPAGE LDS sample buffer (Life Technologies, Carlsbad, CA) and heated at 70°C for 10 min. Equal amount of proteins were loaded onto a 4-12% gradient polyacrylamide gel (Life Technologies) and separated by electrophoresis and transferred to a PVDF membrane (Immobilon-P, Millipore). The membranes were blocked in 5% non-fat milk dissolved in Tris buffered saline with 0.05% Tween-20 (TBS-T) for 1 h at room temperature before probing with primary antibodies overnight at 4 °C, followed by incubation in the appropriate secondary antibody for 1 h at room temperature. The resulting signals were developed using an Enhanced Chemiluminescence (ECL) Western blotting detection reagent kit (GE Health Care, Piscataway, NJ). Digital records were obtained from each blot and the protein bands of interest were quantified using 1-D analysis software (Gel Logic, Carestream, Nutley, NJ). The membranes were stripped and reprobed for GAPDH as the loading control.

### Immunofluorescence microscopy

Fibroblasts were grown in 35 mm cell culture dishes for immunofluorescence assays. For staining of α-SMA, the cells were fixed and permeabilized in a cold acetone:methanol mixture (1:1) for 10 min. For staining of other targeted proteins, the cells were fixed in 4% paraformaldehyde/PBS for 20 min and permeabilized with 0.1% Triton-X-100/PBS for 3 min. After fixation and permeabilization, cells were blocked in 2% goat serum plus 1% Bovine Serum Albumin (BSA) in PBS for 30 min at room temperature, incubated with primary antibody at 4°C overnight, then incubated with secondary antibody conjugated with Alexa Fluor 568 or 488 (Invitrogen/Life Technologies) at room temperature for 2 h. Nuclei were counter-stained with 4,6-diamidino-2-phenylindole (DAPI). Primary antibodies were anti-α-smooth muscle actin (Cat# ab5694, Abcam), anti-MRTF/Mkl1 (Cat# ab49311, Abcam), and anti-SRF (Cat# 5147, Cell Signaling Technology). Filamentous actin (F-actin) was visualized using phalloidin conjugated with Alexa Fluor 488 (Cat# A12379, Invitrogen/Thermo Scientific). Globular monomeric actin (G-actin) was labeled by DNAse I conjugated with Alexa Fluor 594. Images were acquired using a Leica DM5500B upright microscope (Leica Microsystems, GmbH, Wetzlar, Germany) with a Retiga SRV Cooled CCD camera (QImaging, Surrey, BC Canada) and ImagePro Plus software (Media Cybernetics, Rockville, MD, USA).

### Quantitation of HA by Fluorophore-assisted Carbohydrate Electrophoresis (FACE)

The HA content in the cell layer and conditioned media of mouse skin fibroblast cultures was measured as described previously (38). Briefly, after proteolytic digestion and ethanol precipitation, HA in the samples was digested down to disaccharides using hyaluronidase SD (2.5 milliunits/μl; 100741-1A, Seikagaku America, Inc.) and labeled with 2-aminoacridone (Invitrogen) at 6.25 mM in 42.5% Me2SO, 7.5% glacial acetic acid, and 0.625 M sodium cyanoborohydride (1.25 μl/cm^2^ of tissue culture surface area). The labeled HA disaccharides were electrophoresed in a Bio-Rad mini-PROTEAN Tetra system using a gel composition of 20% acrylamide (37.5:1; Bio-Rad), 40 mM Tris acetate (pH 7.0), 2.5 % glycerol, 10% ammonium persulfate, and 0.1% TEMED. After electrophoresis at 500 V (constant voltage) for 50 min at 4 °C, gels were imaged on a UV transilluminator at 365 nm using a CCD camera. The HA disaccharide band was quantified using Gel-Pro Analyzer^®^ version 3.0 (Media Cybernetics, Silver Spring, MD).

### Statistical Analysis

Statistical analyses were performed using a 2-sided Student t-test. *P*<0.05 was considered statistically significant.

## ACKNOWLEDGEMENTS

We sincerely thank Ms. Valbona Cali and Dr. Ronald Midura for their assistance in generating the FACE data, and Dr. Judy Drazba and her team in the Lerner Research Institute’s Digital Imaging Core for assistance with cell imaging. This work was supported by NIH grants P01 HL107147, and S10OD019972.

---

**Supplementary Figure 1.**
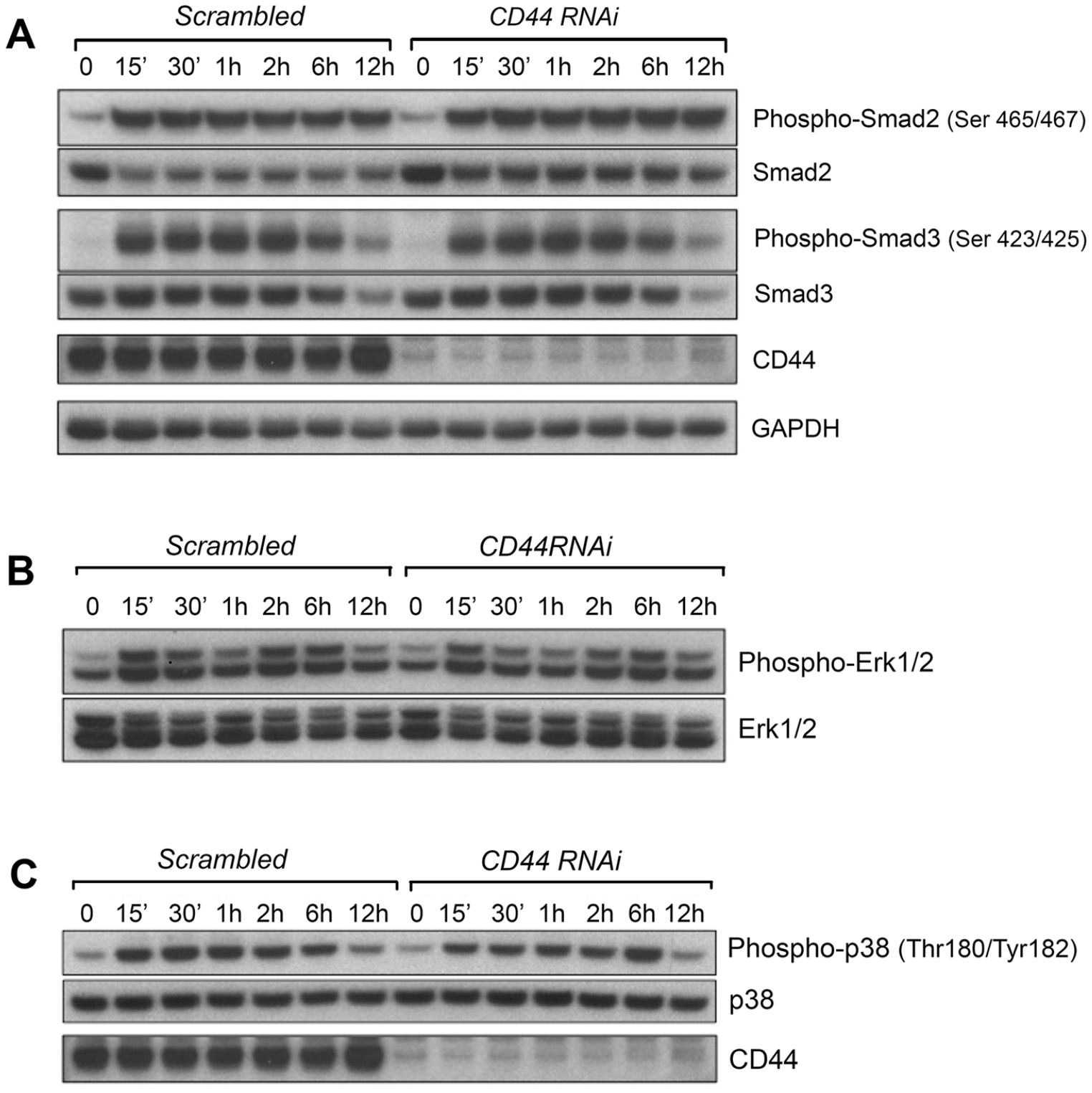
CD44 RNAi has minimal effects upon canonical or non-canonical TGF-β signaling. Primary murine skin fibroblasts were transfected with either scrambled or CD44 RNAi for 48 h, followed by serum starvation for another 24 h to synchronize the cells. Then the cells were re-fed with fresh DMEM media (with 2% FBS) containing 2 ng/mL of rTGF-β1, then harvested at various time points (0, 15 min, 30 min, and 1, 2, 6, and 12 h) after refeeding. (**A**), western blots of Smad2 phosphorylated at residues Serine465/467, and Smad3 phosphorylated at residues Serine 423/425, along with blots of total Smad2 and Smad3. CD44 knockdown efficacy was validated by western blotting of CD44 protein abundance (data not shown). GAPDH was reprobed as a loading control. (**B**), blots of phosphorylated ERK1/2 (at residues Thr202/Tyr204 and Thr185/Tyr187, respectively) and total ERK1/2. (**C**), blots of p38MAPK phosphorylated at residues Threonine180/Tyrosine182, and total p38MAPK. Each western blot is representative of two independent experiments that gave similar results.

**Supplementary Figure 2.**
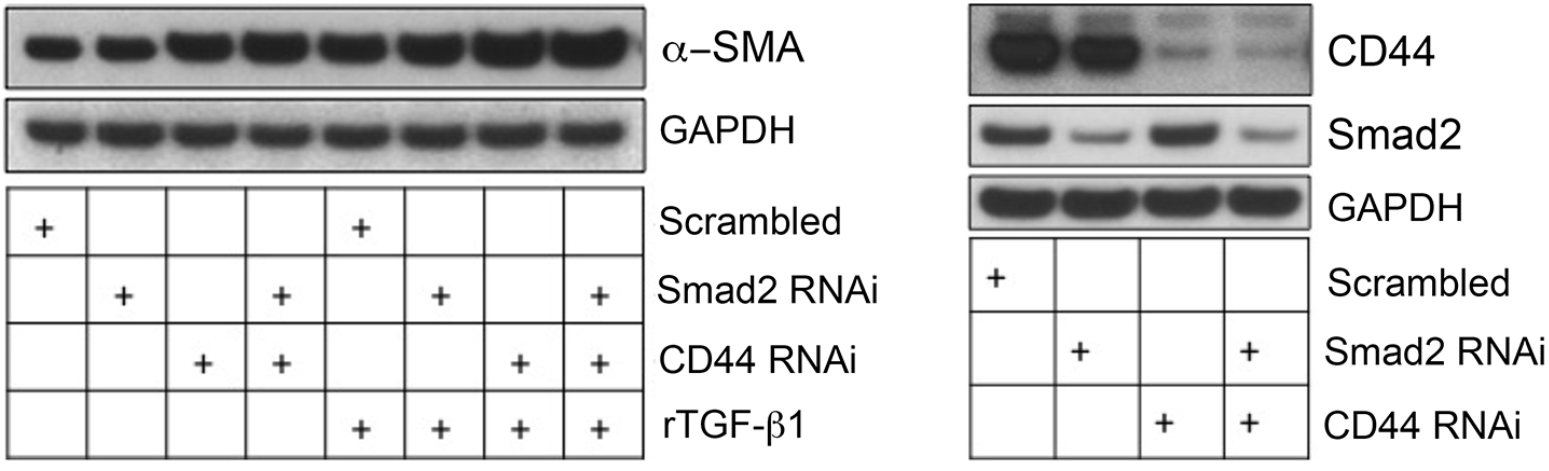
Smad2 RNAi has no effects upon α-SMA gene expression in mouse skin fibroblasts. Fibroblasts were transfected with *Smad2*, *CD44*, or scrambled RNAi, ± rTGF-β1 treatment for 48 h. The protein abundance of α-SMA was examined by western blot with GAPDH as a loading control (*left panel*). Smad2 and CD44 knockdown efficiencies were validated by western blot (*right panel*). Results are representative of 2 experiments.

